# S6K1 and S6K2 regulate homologous recombination DNA repair through control of BRCA1 protein stability

**DOI:** 10.64898/2026.04.28.721439

**Authors:** Mariana Marcela Gois, Laís Bonafé, Luiz Guilherme Salvino da Silva, Mariana Camargo Silva Mancini, Rosalie A. Kampen, Isadora Carolina Betim Pavan, Matheus Brandemarte Severino, Nathalia Quintero-Ruiz, Sylvie M. Noordermeer, Fernando Moreira Simabuco

## Abstract

Recent studies have suggested that S6 kinase 1 (S6K1) contributes to DNA repair (DR). However, the specific pathways and mechanisms involved in this regulation remain unclear. Moreover, it has not been investigated whether S6K2, a functional homologue of S6K1, also contributes to DR. In this study, we investigated the function of both S6K1 and S6K2 (S6K1/2) proteins in DR and demonstrate that both are important for efficient Homologous Recombination-mediated repair (HR). Double knockout of S6K1/2 prevented the formation of BRCA1 and RAD51 foci and increases sensitivity to DNA-damaging agents such as PARP1 inhibitors, cisplatin, and X-ray irradiation. In addition, double knockout of S6K1/2 increased markers of genomic instability, while single knockout had little effect on HR markers and genome stability, which suggests that one kinase can compensate for the loss of the other. Mechanistically, we show that S6K1/2 regulate BRCA1 protein stability, limiting its degradation by the proteasome. Finally, pharmacological inhibition of S6K1/2 sensitised HR-proficient breast cancer cells to Olaparib. Our findings clarify the role of S6K1/2 proteins in HR and suggest that targeting these kinases may be a therapeutic strategy to enhance PARP inhibitor efficacy in HR-proficient tumours.

## INTRODUCTION

The S6K1 and S6K2 (S6K1/2) proteins are the central effector proteins of the mTOR pathway with canonical roles in cell growth, metabolism, and protein translation. S6K1/2 are homologues and exhibit high structural similarity, particularly in their kinase domain (Gout *et al*, 1998), and are known to share certain functions and substrates. For example, both phosphorylate the ribosomal protein S6, their main common target (Tavares *et al*, 2015; Magnuson *et al*, 2012).

Previous research has linked S6K1/2 overexpression and hyperphosphorylation to cancer, especially breast cancer (Filonenko *et al*, 2004). The *RPS6KB1* gene, which encodes the S6K1 protein, is located on chromosome 17q23, a region identified as a common amplification in breast cancer and suggested to be related to the increased expression of genes that favour tumour formation and progression (Sinclair *et al*, 2003). Increased phosphorylation of S6K1 is also associated with the development of resistance to chemotherapy and radiotherapy (Choi *et al*, 2020).

Studies have indicated that S6K1 may participate in DNA repair (Amar-Schwartz *et al*, 2022; Calderon-Aparicio *et al*, 2024). However, the specific pathway and the molecular mechanism regulated by S6K1 remain poorly understood. Moreover, all published studies to date have exclusively analysed S6K1, while S6K2 has been widely neglected. Given their structural similarity, S6K2 might also play unexplored roles in the regulation of DNA repair.

Double-strand breaks (DSB) are the most severe forms of DNA damage, as they generate genomic instability, leading to mutations and potentially to cancer development and cell death (Jackson & Bartek, 2009). The two major DSB repair pathways are Non-Homologous End Joining (NHEJ) and Homologous Recombination (HR)., HR is a high-fidelity DNA repair pathway that that uses the DNA from the sister chromatid as a template to repair DSB without introducing mutations (Takata *et al*, 1998; Jasin & Rothstein, 2013; Scully *et al*, 2019). BRCA1 plays a role in the initial phases of HR, stimulating DNA end-resection, and in the recruitment of RAD51, which, through interaction with PALB2 and BRCA2, searches for the homologous sequence to be used as the template for repair (Zhao *et al*, 2017; Cousineau *et al*, 2005; Ceppi *et al*, 2024).

Deficiencies in HR lead to genomic instability and an increase in mutations that can result in tumour formation. Despite their increased tumour incidence, cells with mutations in HR factors, such as BRCA1, are more sensitive to treatment with DNA-damaging agents, such as cisplatin and PARP inhibitors. The synthetic lethality of PARP inhibition in HR-deficient tumours has revolutionised the personalised treatment of breast and ovarian cancer with BRCA1/2 mutations (Farmer *et al*, 2005; Bryant *et al*, 2005). However, tumours frequently develop resistance to treatment with PARP inhibitors, often due to the re-establishment of HR function. Therefore, continued exploration of the mechanisms of HR function is necessary and potentially important for cancer therapy (Gogola *et al*, 2019).

In this study, we sought to clarify whether S6K1/2 directly regulate DNA repair through Non-Homologous End Joining (NHEJ) and Homologous Recombination (HR), the two major double-strand break repair pathways. We demonstrate that both S6K1/2 influence the functioning of HR, while no effect on NHEJ is observed. Using different genetic models, we demonstrate that S6K1/2 control BRCA1 protein stability and the formation of BRCA1 and RAD51 foci. We also showed that S6K1/2 are functionally redundant, and that in the absence of one S6K, the other can compensate for its activity, requiring a double knockout (DKO) of both S6K1/2 to generate HR defects. Consistent with this, cells with S6K1/2 DKO had increased genomic instability and sensitivity to DNA-damaging agents, such as cisplatin, the PARP inhibitor olaparib and irradiation. Finally, we show that pharmacological inhibition of S6K1/2 sensitised HR-proficient breast cancer cells to PARP1 inhibitors, highlighting a potential therapeutic strategy to enhance PARP inhibitor efficacy in HR-proficient tumours.

## RESULTS

### S6K1 and S6K2 knockdown impairs Homologous Recombination (HR) repair efficiency

To investigate whether the S6K proteins play a role in DSB repair, we used the DSB-Spectrum V1 assay (van de Kooij *et al*, 2022). This HEK293-based fluorescent reporter generates a Cas9-dependent DSB and allows simultaneous analysis of repair via Non-Homologous End Joining (NHEJ) and HR. We combined this system with siRNAs for S6K1 and S6K2 and measured the impact of the knockdown on the efficiency of the repair by the two pathways, using acquired BFP expression as a readout for NHEJ, and GFP as a readout for HR.

The assay was validated using siRNAs against 53BP1 and BRCA1 as controls for NHEJ HR, respectively. The knockdown of S6K1 or S6K2 did not affect the efficiency of NHEJ (Figure 1A, C). In contrast, both siRNAs induced a 50% reduction in HR efficiency. Notably, the knockdown of both S6K1/2 at the same time did not cause a cumulative effect (Figure 1B, C). This suggests that the two proteins play a redundant role in DSB-repair via HR. Alternatively, the limited efficiency of the siRNAs did not allow further reduction, as it was still possible to detect low levels of phosphorylated S6 after the combined knockdown (Figure 1C).

**Figure 1.**
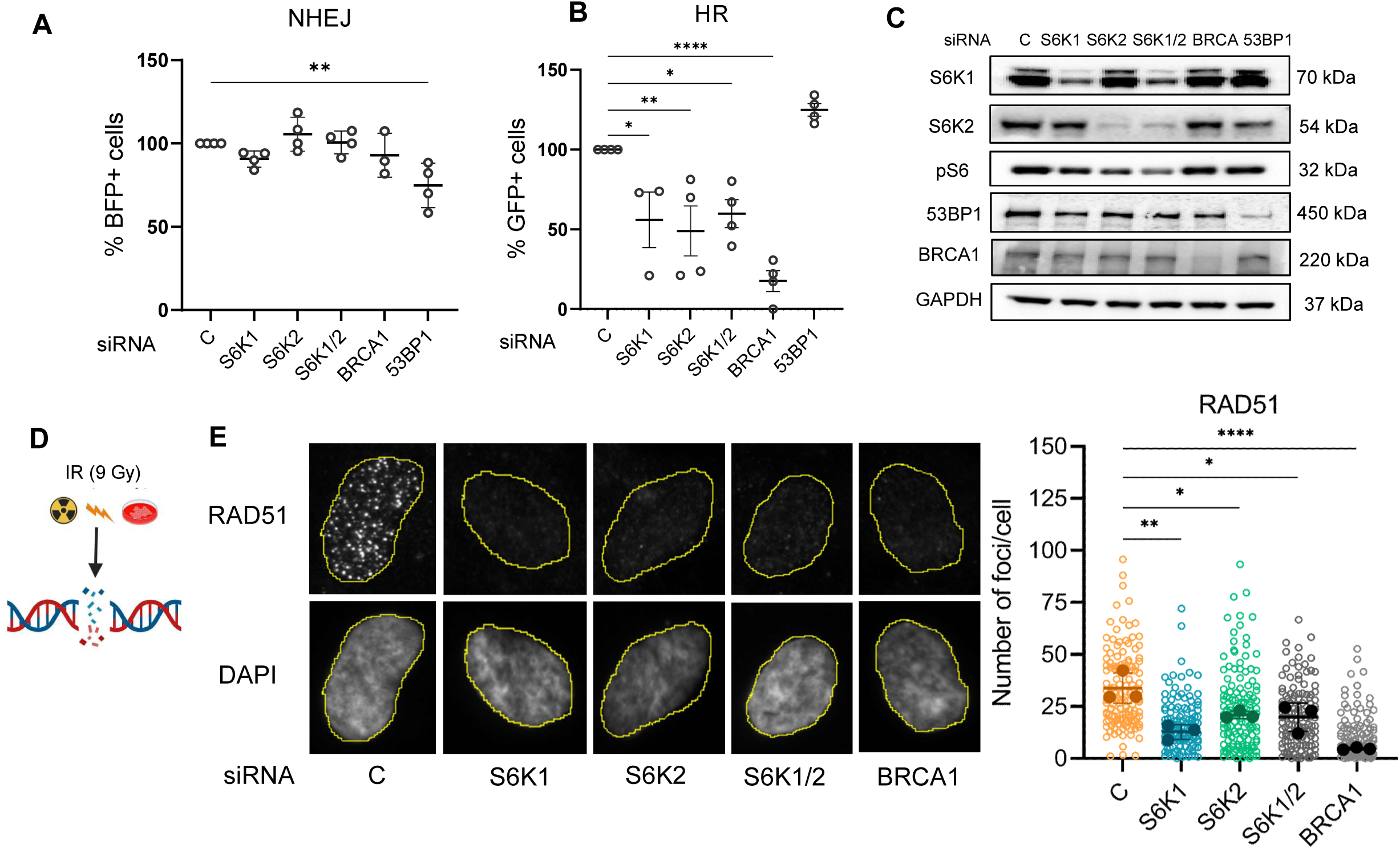
S6K1 and S6K2 knockdown impairs Homologous Recombination (HR) repair efficiency. **A)** and **B)** Cell cytometry results from DSB-Spectrum V1 assay. The percentage of BFP+ or GFP+ cells represents cells positive for repair via NHEJ or HR, respectively. One-way ANOVA with Dunnett’s post-hoc test was used for statistical analysis. Data is presented as mean ± SD of 4 biological replicates. **C)** Western blotting of HEK293 DSB-Spectrum cells subjected to knockdown with siRNAs. GAPDH was used as a loading control. **D)** Scheme of hTERT-RPE1 *TP53* KO cells subjected to X-ray after knockdown of S6K1/2 and BRCA1 for 72 hours. 3 hours post 9 Gy IR, cells were fixed and stained with RAD51 antibody for immunofluorescence analysis. **E)** RAD51 foci were counted using ImageJ. Data represent mean ± SD of three independent biological replicates. Statistical analysis was performed using one-way ANOVA with Dunnett’s multiple comparison test. Data is presented as foci per cell of 3 biological replicates with mean ± SD indicated. *=p<0.05, **=p<0.01, ****=p<0.0001. ≥ 100 cells per group were analyzed in each biological replicate for all experiments

To further validate the role of S6K1/2 in HR, we analysed RAD51 foci formation in hTERT-RPE1 *TP53*-KO cells (van de Kooij *et al*, 2024) upon siRNA transfection against S6K1 or S6K2 or both, and 9 Gy irradiation to induce DSBs (Figure 1D). The knockdown of S6K1/2 alone or in combination decreased the RAD51 foci formation after DNA damage by IR, consistent with the defect in HR found in the reporter assay. Knockdown of the positive control BRCA1 also decreased RAD51 foci formation, as expected (Cousineau *et al*, 2005) (Figure 1E).

### Double-knockout of S6K1/2 impairs BRCA1 and RAD51 foci formation and increases genomic instability

To further investigate the role of S6K1/2 in HR, we used the hTERT-RPE1 *TP53*-knockout (KO) cell line and generated single knockout (SKO) clones of S6K1 or S6K2, and double knockout (DKO) of S6K1/2 simultaneously. We chose a *TP53*-KO cell line to prevent p53-mediated DNA damage responses, including cell cycle arrest and modulation of genes involved in repair (Williams & Schumacher, 2016). This enabled a more specific analysis of the effects of S6K1/2 modulation, without p53-dependent secondary effects (Figure 2A). After delivery of guide RNAs and Cas9 and expansion of single cells, the genotypes were confirmed by Sanger sequencing (Figure S1). For subsequent experiments, we used two independent DKO clones (#30.7 and #30.11), two SKO clones for S6K1 (#30 and #32), and two SKO clones for S6K2 (#16 and #18). As expected, due to redundant functions of S6K1/2 (PENDE et al., 2004), only DKO cells could abolish the phosphorylation of S6 to near completion (Figure 2B).

**Figure 2.**
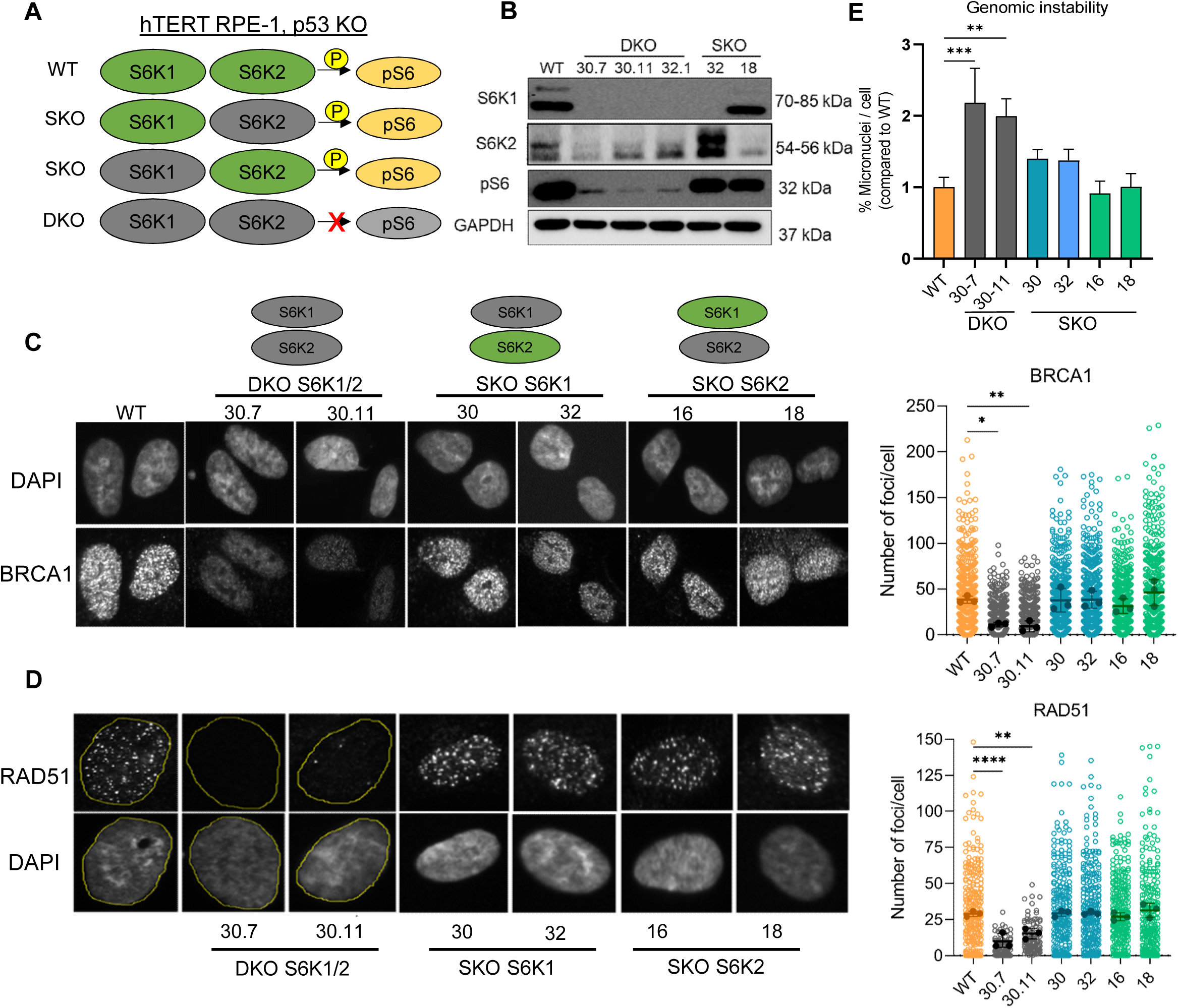
Double-knockout of S6K1/2 impairs RAD51 and BRCA1 foci formation and increases genomic instability. **A)** Scheme of hTERT-RPE-1 cells edited with single and double-knockouts (SKO and DKO) of S6K1 and S6K2, and their phenotype regarding S6 phosphorylation. **B)** Western blotting of SKO and DKO clones. **C)** and **D)** WT, SKO, and DKO cells were treated with 9 Gy irradiation. 3 hours post IR, cells were fixed and stained with DAPI, RAD51, and BRCA1 antibodies for immunofluorescence analysis. Foci were counted using ImageJ. One-way ANOVA with Dunnett’s post-hoc test was used for statistical analysis. Results are presented as foci per cell of 3 biological replicates with mean ± SD indicated. ****=p<0.0001. **E)** Genomic instability in RPE-1 cells was quantified by the number of micronuclei per cell after 9 Gy IR. Three hours post IR, cells were stained with DAPI, and the number of micronuclei was counted manually. The percentage of micronuclei in DKO and SKO cells was calculated and compared with WT cells. One-way ANOVA with Dunnett’s post-hoc test was used for statistical analysis. Data is presented as mean ± SD of 3 biological replicates. **=p<0.01 and ***=p<0.001. ≥ 100 cells per group were analysed in each biological replicate for all experiments

We exposed RPE1 wild-type (WT) cells, as well as SKO and DKO clones of S6K1/2, to 9 Gy of IR and analysed BRCA1 and RAD51 foci formation 3 hours post IR. DKO clones showed a marked reduction in both BRCA1 and RAD51 foci after IR. In contrast to our data using siRNAs, SKO clones showed levels of BRCA1 and RAD51 foci comparable to WT cells (Figures 2C and 2D).

This suggests that in a chronic knock-out setup – in contrast to acute depletion - cells adapt and compensate for the loss of one S6K through the activity of the other, similarly to their redundant role in S6 phosphorylation, also validated in our models (Figure 2B) (Pende *et al*, 2004).

Since defects in DNA repair can promote genomic instability, we next analysed whether our models showed genome instability markers by quantifying micronuclei formation (MacDonald *et al*, 2024). Consistent with the observed HR defects, DKO cells exhibited a two-fold increase in micronuclei compared to WT cells, whereas SKO cells showed levels comparable to WT (Figure 2E).

### Double-knockout of S6K1 and S6K2 sensitize cells to DNA damage

Next, we analysed the sensitivity of our models to DNA-damaging drugs. DKO cells showed increased sensitivity to olaparib (PARP1 inhibitor) and cisplatin when compared to WT cells. In contrast, SKO cells showed similar sensitivity as WT cells (Figure 3A-E). We also analysed the sensitivity to 7 Gy of X-ray irradiation using clonogenic survival assays. The DKO cells markedly formed fewer colonies than WT cells, and SKO cells formed colonies comparable to WT cells (Figure 3F). All together, our results suggest that S6K1/2 control the repair of DSB via HR. Moreover, the resistance of SKO cells to DNA-damaging agents supports our hypothesis that S6K1 and S6K2 function redundantly in HR, with one kinase compensating for the loss of the other in the SKO models.

**Figure 3.**
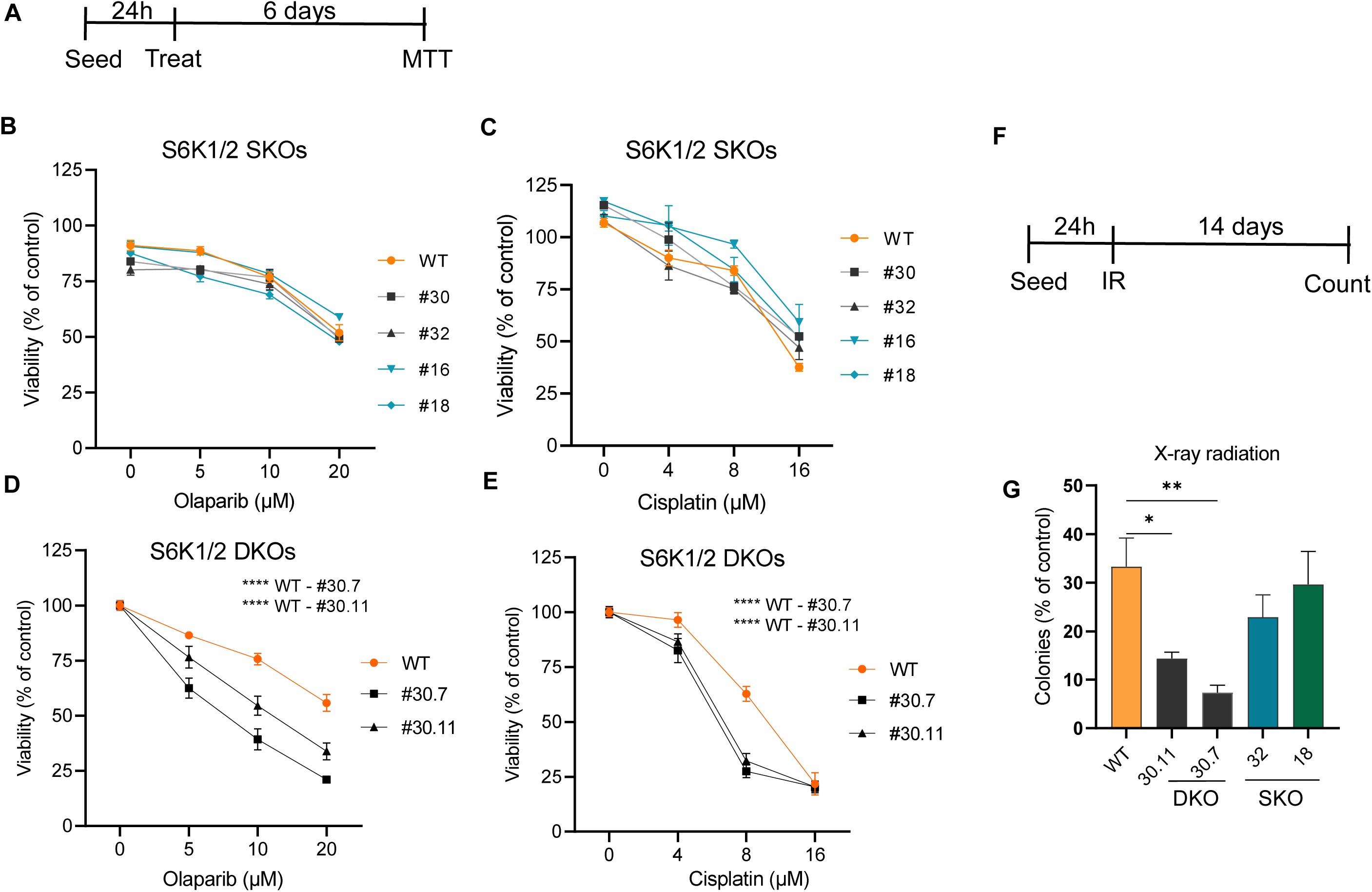
Double knockout of S6K1/2 sensitises cells to DNA damage. **A)** Scheme used for MTT assay. hTERT-RPE-1 WT, SKO and DKO clones were treated with olaparib or cisplatin for 6 days and then subjected to MTT assay for viability analysis. Each well had the absorbance measured at 570 nM, and the drug-treated conditions were normalized to the control treated with the vehicle. WT and SKO results are represented in **B)** and **C)**. WT and DKO results are represented in **D)** and **E)**. Two-way ANOVA with Dunnett’s post-hoc test was used for statistical analysis, and data are presented as mean ± SD of 3 biological replicates for B-E. **F)** Scheme used for clonogenic survival assay. **G)** RPE-1 WT, SKO, and DKO cells were seeded at low density and treated with 7 Gy of IR. After 14 days, the colonies were stained with crystal violet and manually counted. The number of colonies was normalized to the non-treated cells. One-way ANOVA with Dunnett’s post-hoc test was used for statistical analysis, and data are presented as mean ± SD of 3 biological replicates. *=p<0.05, **=p<0.01, ***=p<0.001, ****=p<0.0001.

### S6K1/2 regulate HR through the regulation of BRCA1 protein stability

To further investigate the mechanism by which S6K1/2 regulate HR, we analysed the protein levels of BRCA1. We found that the DKO cells have reduced BRCA1 protein levels, while SKO cells have BRCA1 protein levels similar to WT cells (Figure 4A). Next, we evaluated gene expression and found that both SKO and DKO cells express BRCA1 mRNA similar to WT cells (Figure 4B). This suggests that the regulation of BRCA1 levels by S6K1/2 is post-transcriptional. Then, we treated both WT and DKO cells with the proteasome inhibitor MG132, and found that MG132 treatment rescued BRCA1 levels in DKO cells (Figure 4C). These data suggest that S6K1/2 regulate BRCA1 stability by limiting its degradation via the proteasome. Moreover, we found that DKO cells exhibit increased DNA damage, evidenced by elevated γH2AX levels, validating their HR deficiency (Racca *et al*, 2021; Guo *et al*, 2025; Zámborszky *et al*, 2017) (Figure 4C).

**Figure 4.**
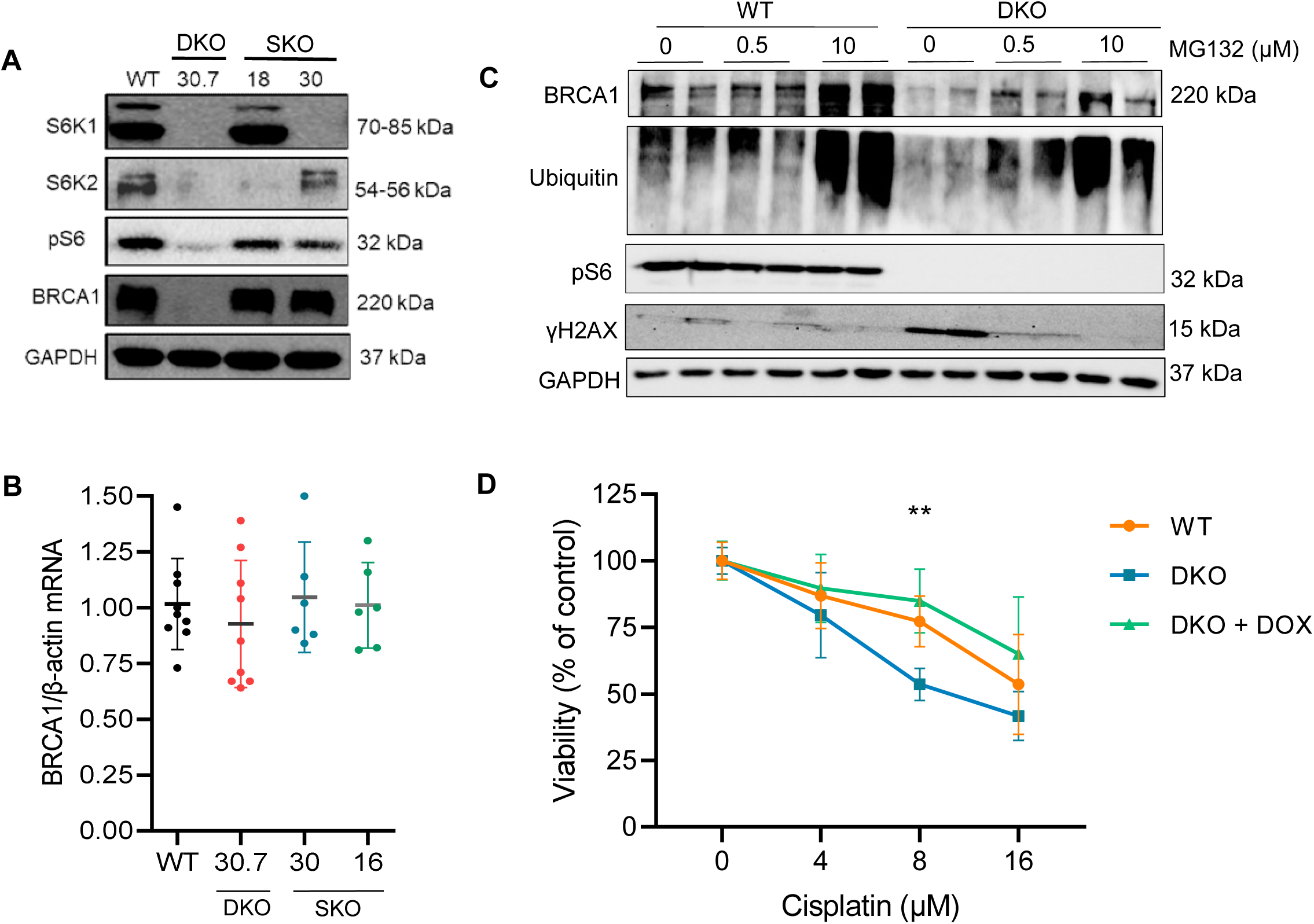
S6K1/2 regulate HR through the regulation of BRCA1 protein stability. **A)** Basal protein levels of hTERT-RPE-1 WT, SKO and DKO clones. **B)** BRCA1 mRNA of RPE1 WT, DKO, and SKO cells was analysed by RTqPCR. β-actin was used as a housekeeping control for normalization. One-way ANOVA with Dunnett’s post-hoc test was used for statistical analysis. Data are presented as mean ± SD of 3 biological replicates. **C)** WT and DKO cells were treated with MG132, and protein levels were assessed by Western blotting. **D)** WT and DKO cells were transduced with a doxycycline(dox)-inducible BRCA1 expression vector. Cells with or without dox (100 ng/mL) were treated with cisplatin for 5 days then MTT assay was performed. Two-way ANOVA with Dunnett’s post-hoc test was used for statistical analysis, and results are presented as mean ± SD of 3 biological replicates. **=p<0.01.

Next, we transduced both WT and DKO cells with a doxycycline (dox)-inducible BRCA1 expression vector (Figure S2) and evaluated whether BRCA1 re-expression could rescue the sensitivity to DNA damaging agents in the DKO cells. Indeed, we found that dox treatment in the complemented DKO mitigated the increased sensitivity to cisplatin (Figure 4D). These data suggest that increased sensitivity to cisplatin of S6K1/2 deficiency is related to a downregulation of BRCA1.

### S6K inhibitors sensitise breast cancer cells to PARPi

Given the critical role of the HR pathway in determining the response of cancer cells to PARP1 inhibitors, we investigated whether inhibiting S6K1/2 could influence the sensitivity of breast cancer cells to olaparib. To this end, we used the cell lines MDA-MB-231, HCC1937, and MCF7, which differ in their characteristics and BRCA1 status but are all considered HR-proficient (Meijer *et al*, 2024). Cells were treated with olaparib alone or after pre-treatment with the S6K1 inhibitor PF4708671 (PF47). PF47 was used in a concentration that is known to inhibit both S6K1 and S6K2 (Pearce *et al*, 2010). Notably, pre-treatment with PF47 significantly increased sensitivity to olaparib in all breast cancer cell lines analysed, compared with cells treated with olaparib alone (Figure 5A–D).

**Figure 5.**
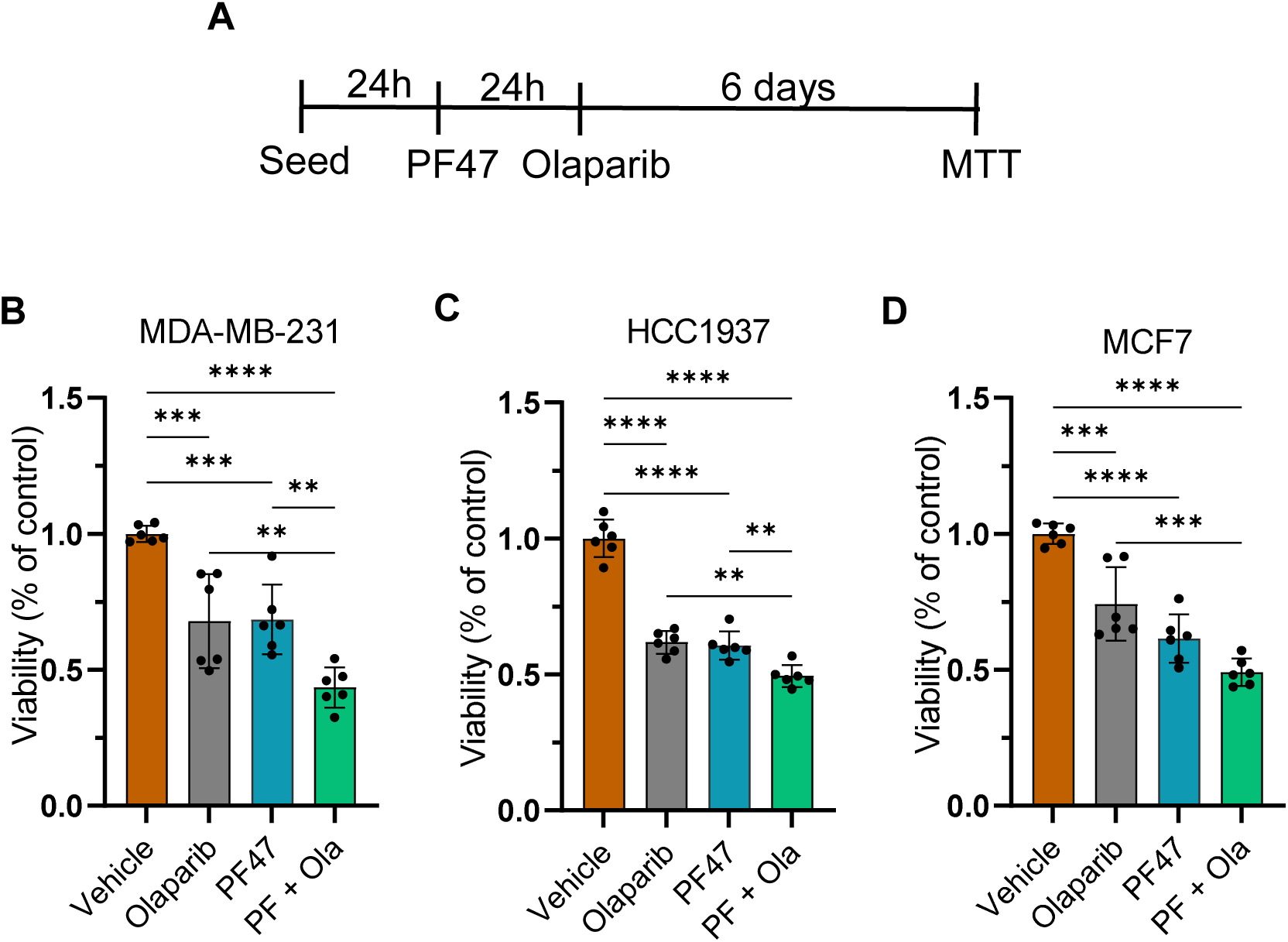
S6K1 inhibitor sensitizes breast cancer cells to olaparib. **A)** Scheme of treatment used in the experiment. **B), C)** and **D)** Treatment of MDA-MB-231, HCC1937, and MCF7 cells with Olaparib and PF4708671 (PF47) alone or with PF47 pretreatment followed by olaparib treatment (PF + Ola). The PF47 dose used for all cells was 20 µM for 24 hours. The olaparib dose used for MDA-MB-231 and HCC1937 was 80 µM, and for MCF7 it was 160 µM for 6 days. Two-way ANOVA with Dunnett’s post-hoc test was used for statistical analysis and results are presented as mean ± SD of 3 biological replicates. **=p<0.01; ***=p<00.1; ****=p<000.1.

## DISCUSSION

The mTOR pathway is traditionally associated with cell growth, metabolism, and protein translation, but accumulating evidence also implicates it in the DNA damage response and repair (Chen *et al*, 2010; Mattoo *et al*, 2019; Danesh Pazhooh *et al*, 2021). However, the specific role of the mTOR effector kinases S6K1 and S6K2 (S6K1/2) in this process remains unclear.

S6K1 has been reported to regulate DNA mismatch repair by phosphorylating MSH6 and indirectly regulate HR repair by phosphorylating CDK1 (Amar-Schwartz *et al*, 2022). Its depletion has also been linked to increased radiation sensitivity of lung cancer cells, possibly by modulating the MRN complex (Wang *et al*, 2017; Calderon-Aparicio *et al*, 2024). Our group reported that S6K2, but not S6K1, interacts with proteins of the NHEJ pathway, including PARP1 and Ku70 (Pavan *et al*, 2016). Considering these observations, which DNA repair pathway is regulated by S6K1/2, and the precise molecular mechanism underlying this regulation remains unclear. Moreover, previous studies have focused exclusively on S6K1, and the role of S6K2 in DNA damage repair remains unexplored (Pardo & Seckl, 2013). In this work, we investigated the functions of both S6K1/2 in DNA double-strand break repair and defined their contributions to the DNA damage response.

Depletion of either S6K1 or S6K2, alone or in combination, decreased HR efficiency without affecting NHEJ (Figure 1A–D). These results are consistent with previous studies implicating S6K1 in HR regulation (Calderon-Aparicio *et al*, 2024; Amar-Schwartz *et al*, 2022) and further indicate that S6K2 may also participate in the same pathway. Combined knockdown of S6K1 and S6K2 did not result in an additive reduction in HR, supporting a model in which the two kinases play similar or redundant roles in HR control. A limitation of the DSB-Spectrum assay is that it relies on siRNA efficiency, which appears to be only partially effective in the combined S6K1/2 depletion, as indicated by the residual phosphorylation of the S6 target (Figure 1C).

We explored the redundancy of S6K1/2 and found evidence of compensatory signalling between the two kinases in the regulation of HR markers in full knock-out clones. We found that DKO cells have impaired BRCA1 and RAD51 foci formation after irradiation, while SKO cells have no defects. This result is consistent with previous work that demonstrated that the mTOR inhibitor rapamycin inhibits HR through decreasing RAD51 and BRCA1 foci formation (Chen *et al*, 2010; Mattoo *et al*, 2019).

The characteristic of S6K1/2 compensation has been previously reported in mice, where a compensation in S6 phosphorylation was demonstrated when only one of the S6K1/2 genes was knocked out (H *et al*, 1998; Pende *et al*, 2004). The same phenotype is visible in the clones we generated in this study. Only DKO clones show strongly decreased levels of S6 phosphorylation, while SKO clones are able to maintain S6 phosphorylation levels comparable to WT cells (Figure 2B). These overlapping or redundant functions are characteristics often found in homolog kinases, which frequently have the ability to buffer each other’s loss (De Kegel & Ryan, 2019).

DKO cells displayed markers of genomic instability frequently found in HR-deficient cells, including micronuclei formation (Baert *et al*, 2020; Ban *et al*, 2001; MacDonald *et al*, 2024), increased γH2AX levels and increased sensitivity to DNA-damaging agents (Telli *et al*, 2016; ter Brugge *et al*, 2023; Farmer *et al*, 2005). These findings confirm that S6K1/2 loss generates an HR-deficient phenotype.

Mechanistically, our results indicated that S6K1/2 regulate BRCA1 protein stability at the post-translational level, rather than transcriptionally. This regulation is likely to involve proteasomal degradation, as BRCA1 levels were restored in DKO cells by proteasomal inhibition (Figure 4A-C). This data is consistent with prior work showing that BRCA1 is degraded by the ubiquitin-proteasome system (Blagosklonny *et al*, 1999; Guo *et al*, 2024; Lu *et al*, 2007). Moreover, mTOR inhibition is associated with increased proteasome activity (Zhao *et al*, 2015). Notably, the re-expression of BRCA1 in DKO cells rescued the resistance to cisplatin (Figure 4D), indicating that BRCA1 destabilization underlies the HR defects caused by S6K1/2 loss (Figure 6).

**Figure 6.**
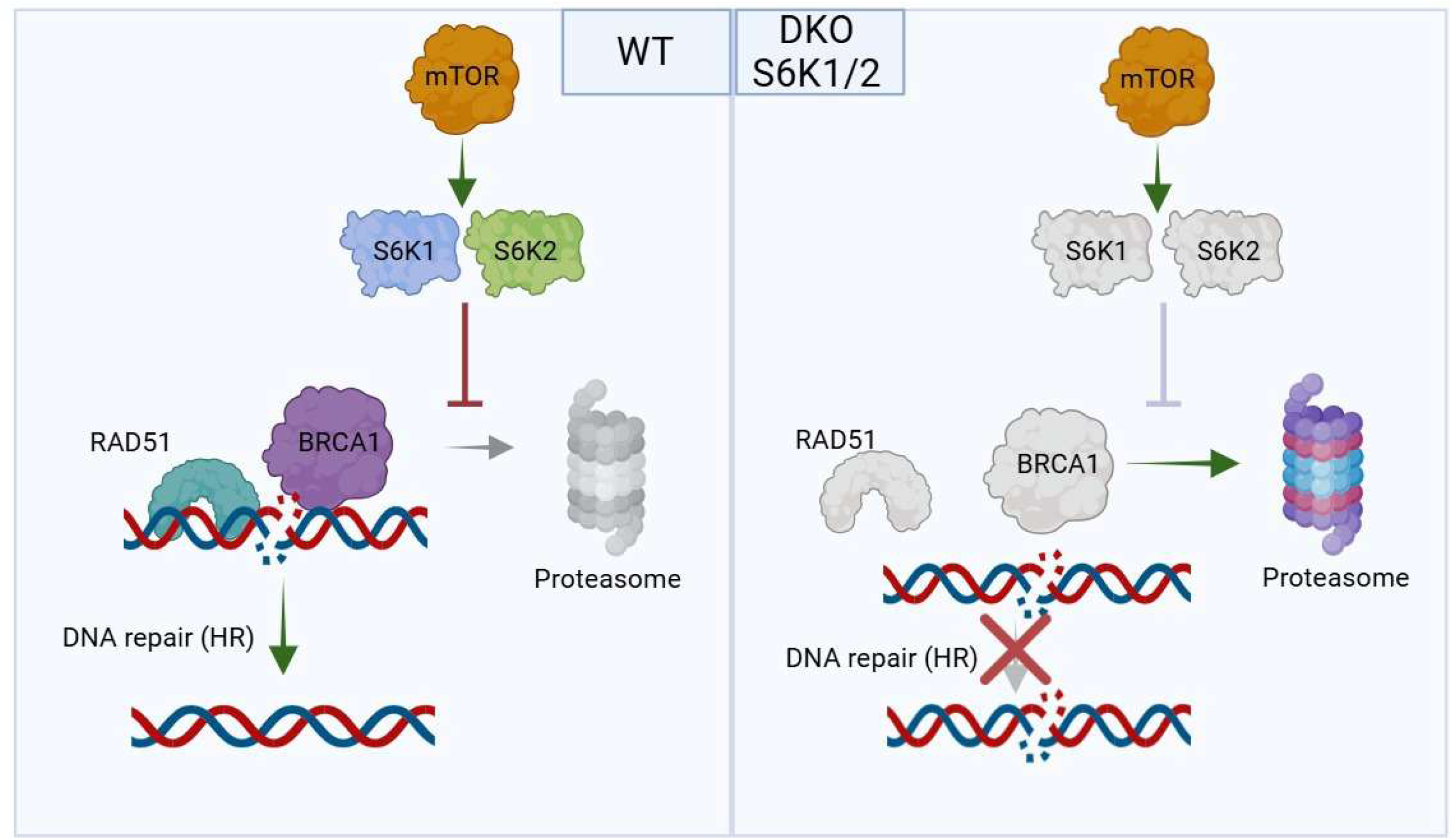
Schematic representation of the proposed model. In wild-type cells, S6K1 and S6K2 maintain BRCA1 stability by limiting its proteasomal degradation, thereby enabling BRCA1 and RAD51 foci formation and efficient homologous recombination (HR). Double knockout of S6K1/2 reduces BRCA1 stability, impairing BRCA1 and RAD51 foci formation, increasing genomic instability and compromising HR.

The expression of the deubiquitinase USP4, a known regulator of BRCA1 stability (Guo *et al*, 2024), showed a strong positive correlation with S6K1 and mTOR (R = 0.73 and 0.79, respectively), and a weak correlation with S6K2 (R = 0.37) (Figure S3). USP4 deubiquitinase activity is controlled by its phosphorylation at Ser445 by the AKT protein (Zhang *et al*, 2012). The Ser445 of USP4 is located in a region within the RXRXXS/T motif, which is recognized by AKT and S6K1/2 (Moritz *et al*, 2010). Moreover, BRCA1 itself also has the same motif at T509 and S694, which are known to mediate BRCA1 protein stability (Nelson *et al*, 2010). Therefore, we raise the hypothesis that S6K1/2 may regulate BRCA1 protein stability through its direct phosphorylation or indirectly through USP4 phosphorylation. Further experiments are needed to conclude which of these mechanisms predominates.

Given the role of S6K1/2 in breast cancer (Filonenko *et al*, 2004; Segatto *et al*, 2013) and the clinical relevance of PARP inhibitors (Lord & Ashworth, 2017), we evaluated this relationship in breast cancer cells. The literature shows synergy between PI3K-AKT-mTOR pathway inhibitors and PARP inhibitors, and several studies have demonstrated that mTOR inhibitors sensitize breast cancer cells to PARP inhibitors (De *et al*, 2014; Li *et al*, 2021; Osoegawa *et al*, 2017; Philip *et al*, 2017).

However, complete mTOR inhibition with rapamycin causes significant side effects that limit its therapeutic use, especially in cancer patients (Bruss *et al*, 2022; Peng *et al*, 2022). The S6K1 inhibitor, PF-4708671 (PF47), has demonstrated antitumor effects in breast cancer cell lines (Choi *et al*, 2013; Hong *et al*, 2013; Khotskaya *et al*, 2014; Segatto *et al*, 2013), but its combination with PARP inhibitors has not been tested.

Our data show that PF47 sensitizes breast cancer cells to olaparib, increasing its cytotoxic effect (Figure 5A-D). Although PF47 is described as S6K1-specific, we used it in a concentration known to cause S6K2 inhibition too (Pearce *et al*, 2010). These findings suggest that S6K1/2 inhibition potentiates PARP1-targeted therapy. We hypothesize that using specific inhibitors such as PF47 in the clinic may generate fewer side effects than global inhibition of the mTOR pathway, while maintaining the benefits of the combination with olaparib. Furthermore, considering that S6K1/2 are involved in the development of resistance to antitumor therapies (Choi *et al*, 2020), their inhibition may represent a possible strategy to increase the sensitivity of resistant tumours.

In summary, we described how S6K1/2 act as key regulators of BRCA1 stability, a previously uncharacterized function of S6K1/2, clarifying the link between the mTORC1–S6K1/2 signalling axis and the maintenance of genome stability through control of homologous recombination DNA repair. Our findings might clarify the mechanism by which S6K1/2 overexpression correlates with chemotherapy and radiotherapy resistance (Choi *et al*, 2020), and suggest that their inhibition may represent a strategy to potentiate PARP inhibitor therapy in HR-proficient tumours.

## MATERIALS AND METHODS

### Plasmids

For CRISPR-Cas9-mediated knockout (KO) of S6K1/2, guide RNAs targeting exon 2 of S6K1 and exon 3 of S6K2 were cloned into pSpCas9(BB)-2A-Puro V2.0 (PX459) (Addgene #62988). For BRCA1 lentiviral expression, the full-length BRCA1 was previously cloned from the pCL-MFG-BRCA1 (Addgene #12341) into pCW57.1_Puro (Addgene #41393) containing a tetracycline-inducible promoter (Tet-On) using gateway cloning, generating the plasmid pCW57.1_Puro_BRCA1-3xTy.

### Cell lines and cell culture

hTERT RPE-1 *TP53*-KO cell line (RPE-1) was previously described (van de Kooij *et al*, 2024) and was used to generate S6K1/2 knockouts. This cell line has a near-diploid stable genome, which makes it suitable for genome integrity studies. HEK293T cells were used to produce the lentivirus. RPE-1 and HEK293T were obtained from American Type Culture Collection (ATCC), and DSB-Spectrum_V1 cells were previously described (van de Kooij *et al*, 2022).

RPE-1, HEK293T and DSB-Spectrum V1 cells were cultured in Dulbecco’s Modified Eagle’s Medium (DMEM) high glucose, GlutaMAX™, and pyruvate supplemented (ThermoFisher Scientific), with 10% Fetal Bovine Serum (FBS). RPE-1 and HEK293T were cultured with 1% penicillin/streptomycin, and DSB-Spectrum V1 cells were cultured without antibiotics.

MCF7, MDA-MB-231 and HCC1937 breast cancer cell lines were originally obtained from the ATCC and were a gift from Prof. Sandra Martha Gomes Dias. All breast cancer cell lines were cultured in Roswell Park Memorial Institute (RPMI) 1640 Medium with 10% Fetal Bovine Serum (FBS) and 1% penicillin/streptomycin antibiotics.

All cells were maintained at 37 °C in a humidified atmosphere containing 5% carbon dioxide and were used for experiments within 2-10 passages from thawing. All cell lines were tested negative for mycoplasma contamination.

### Genetic editing of cells

To obtain clonal S6K1 and S6K2 knockouts, guide RNAs targeting S6K1 or S6K2, previously cloned into pSpCas9(BB)-2A-Puro V2.0 (PX459) plasmid (Addgene #62988) were nucleofected into hTERT RPE-1 *TP53*-KO cell line (hereafter also referred to as Wild-type (WT). Nucleofected cells were then selected with 2 µg/mL of puromycin for 48 hours, and clones were isolated by seeding 500-1000 cells in 15 cm dishes. Single colonies were expanded to 24, 12, and 6-well plates, then genomic DNA was isolated, and the region of interest was amplified and sequenced using Sanger sequencing.

### Viral transductions

Lentiviruses were produced in HEK293T cells by jetPEI transfection (Polyplus) of plasmid pCW57.1_Puro_BRCA1-3xTy with third-generation packaging vectors pMDLg/pRRE, pRSV-Rev, and pMD2.G. Viral supernatants were collected 48-72 hours post-transfection, filtered with 0.45 µm filter, and used to transduce cells in the presence of 4 µg/mL polybrene. For RPE1, 2 µg/mL of puromycin was used to select transduced cells.

### Drugs

The following drugs were used: Olaparib (SelleckChem), Cisplatin (Accord), MG-132 (Sigma), PF-4708671 (Sigma).

siRNA used:

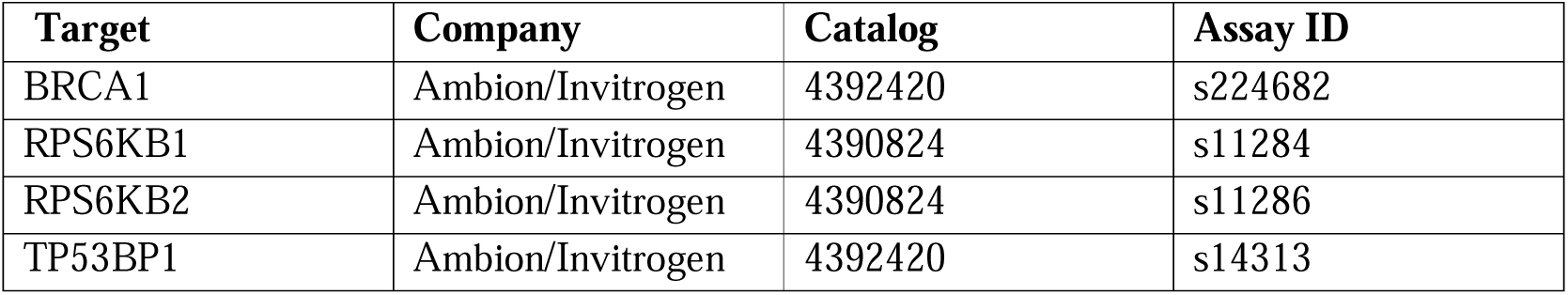

### X-ray radiation

DNA damage was introduced using an XYion X-ray radiation system. First, a radiation dosimeter was used to calculate the amount of Grays (Gy) absorbed per minute at the selected area. Then, the plates with cells were positioned in the same area for the time needed to reach the designated amount of Gy.

### Clonogenic survival assays

Cells were seeded in 10-cm dishes with densities between 500 and 1000 cells; the next day, cells were irradiated with 7 Gy and incubated at 37 °C. After 14 days, colonies were stained with crystal violet solution (0.4% crystal violet, 20% methanol) and manually counted. Relative survival was calculated by setting the number of colonies in non-treated controls at 100%.

### Viability assays (MTT)

For viability assays, cells were seeded in 96-well plates with densities between 800 to 1,000 cells per well. Plates were incubated at 37 °C in the presence of a drug or vehicle (e.g., DMSO). After the treatment, MTT (Sigma) was added for 3 hours, and formazan crystals were solubilized with HCL 1M: isopropanol (1:25) solution for 15 minutes. The absorbance was measured at 570 nM.

### DSB-spectrum_V1 assay

The DSB-Spectrum_V1 reporter assay was performed as previously described (van de Kooij *et al*, 2022). HEK 293T DSB-Spectrum_V1 cells were seeded, and the next day, transfected with siRNAs. After 24 hours, a second siRNA transfection was performed. After 6-8 hours from the second siRNA transfection, 20,000 cells were seeded per well in 96-well plates. After 24 hours from seeding, the cells were transfected in technical triplicate with the pX459-Cas9-sgRNA-iRFP construct containing either sgBFP targeting DSB-Spectrum or sgAAVS1. The cells were analyzed by flow cytometry 48-96 hours after the DSB-Spectrum targeting transfection. FlowJo software (BD Biosciences) was used to analyze the flow cytometry data. Gating on forward and side-scatter was applied to select the live and single-cell population. Transfected cells were selected by gating on iRFP. In this population, frequencies of BFP+ and GFP+ cells were quantified. The frequency of each fluorescent sub-population in the sgAAVS1-transfected cells was subtracted from the frequency of that same population in the sgBFP-transfected cells. The resulting background-corrected frequencies were normalized to the siCTRL-transfected cells.

### IR-induced foci immunofluorescence (IRIF)

For RAD51 and BRCA1 immunofluorescence, cells were grown on glass coverslips until 70-90% confluence and irradiated with 9 Gy of X-rays. After 3 hours, cells were treated with cold nuclear extraction (NuEx) buffer (20 mM Hepes pH 7.5, 20 mM NaCl, 5 mM MgCl_2_, 1 mM DTT, 0.5% NP-40 (Sigma Aldrich), 1× Complete Protease Inhibitor Cocktail (Sigma Aldrich) for 12 minutes at 4°C. Then, cells were fixed with 2% paraformaldehyde in PBS for 20 minutes at room temperature. Next, cells were washed 3 times with PBS and blocked with PBS+ (5 g/L BSA (Sigma Aldrich) and 1.5 g/L glycine (Sigma Aldrich) in PBS) for 30 minutes at room temperature. Cells were incubated for 1.5-2 hours with the primary antibody in PBS+, washed 5 times with PBS, and incubated for 1-1.5 hours with DAPI 0.1 μg/mL and the secondary antibody in PBS+. After washing 3 times with PBS, the coverslips were mounted using Aqua-Poly/mount (Polysciences). Pictures were taken using the Zeiss Axio Imager 2 fluorescent microscope at a 40x zoom. Foci of at least 100 cells per condition per replicate were quantified using the IRIF analysis 3.2 Plugin in ImageJ.

Antibodies used for immunofluorescence:

| **Antibody** | **Dilution** | **Brand** | **Catalog** |
| --- | --- | --- | --- |
| BRCA1 | 1:1000 | Merck Millipore | #OP92 |
| RAD51 | 1:1000 | Abcam | #ab176458 |

### Western-blotting

Protein was extracted using a cell lysis buffer (50 mM Tris-Cl, pH 7.5, 150 mM NaCl, 1 mM EDTA, 1% Triton X-100) supplemented with Complete Protease and Phosphatase Inhibitor Cocktail (Sigma Aldrich). SDS sample buffer with DTT was added to the lysates, followed by denaturation at 95°C for 5 minutes. Total protein content was separated using electrophoresis with NuPAGE Bis-Tris 4-12% polyacrylamide gels (ThermoFisher). Proteins were transferred to nitrocellulose membranes and blocked for 1 hour at room temperature with 5% BSA (Sigma Aldrich) in TBS-T (50 mM Tris-Cl, pH 7.5; 150 mM NaCl; 0.1% Tween-20). Membranes were then incubated overnight at 4°C with appropriate primary antibodies, and then incubated for 1-1.5 hours at room temperature with HRP-labelled secondary antibodies. Western Bright ECL HRP Substrate kit (Advansta) was used to generate chemiluminescence. Images of membranes were taken using the Chemidoc Imaging Systems (Biorad).

Antibodies used for Western-blotting:

| **Antibody** | **Dilution** | **Brand** | **Catalog** |
| --- | --- | --- | --- |
| BRCA1 | 1:1000 | Cell Signaling | #14823 |
| γH2AX | 1:1000 | Cell Signaling | #9718 |
| phospho-S6-Ser240/244 | 1:2000 | Cell Signaling | #5364 |
| GAPDH | 1:2000 | Cell Signaling | #2118 |
| S6K1 | 1:1000 | Cell Signaling | #2708 |
| S6K2 | 1:1000 | Cell Signaling | #14130 |
| 53BP1 | 1:2000 | BD Biosciences | #612522 |
| Ubiquitin | 1:1000 | Cell Signaling | #3933 |

### RTqPCR

The cells were plated in a 6-well plate at a density of 1×10^5^ cells. Total RNA was extracted using TRIzol (Thermo Scientific). The High-Capacity cDNA Reverse transcription kit (Thermo Scientific) was used to synthesize the cDNA. iTaq Universal SYBR Green Super Mix (Bio-Rad) was used following the manufacturer’s instructions. The Gene expression was analyzed by the formula: 2^-ΔΔCt^ (Livak & Schmittgen, 2001), using β-actin as a housekeeping gene. Samples were arranged in triplicate in a 96-well plate for amplification and were run in the Step One Plus Real-Time PCR System (Applied Biosystems).

Primers used for RTqPCR:

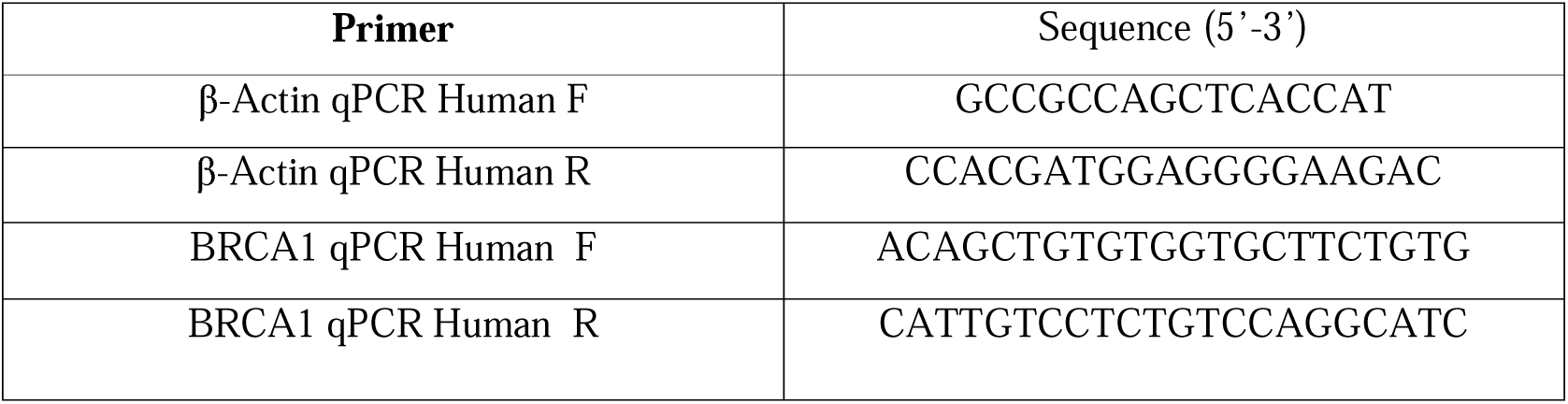

### Bioinformatic analysis

Benchling web server (https://www.benchling.com/) was used to visualize DNA sequences. GEPIA was used for correlation analysis (http://gepia.cancer-pku.cn/). Inference of CRISPR Edits (ICE) analysis was used for decomposition of Sanger sequencing data to characterize the genotype of CRISPR-edited clones (https://ice.editco.bio/).

### Statistical analysis

Statistical analyses were performed using GraphPad Prism (version 9.1.0). Data are presented as mean ± SD or 95% confidence interval, as specified in the figure legends. For experiments involving two independent variables, two-way ANOVA followed by Dunnett’s post-hoc test was used to compare conditions to the respective control. For single-variable comparisons, one-way ANOVA with Dunnett’s post-hoc test was applied. At least 3 independent biological replicates were repeated for each experiment. Statistical significance was defined as p < 0.05.

## FUNDING INFORMATION

This study was supported by the São Paulo Research Foundation (FAPESP), grant numbers 2018/14818-9 (FMS), 2023/02842-0 (MMG), 2020/08684-0 (MMG) and by Oncode Institute (SMN).

## AUTHOR CONTRIBUTION

MMG: conceptualization; methodology; data curation; formal analysis; investigation; visualisation; writing - original draft, review and editing; funding acquisition. LMCBO: data curation; investigation; writing - review and editing. LGSS: data curation; investigation; writing - review and editing. MCSM: data curation; investigation; writing - review and editing. RAK: data curation; investigation; writing - review and editing. ICBP: data curation; investigation; writing - review and editing. MBS: data curation; investigation; writing - review and editing. NQR: data curation; investigation; writing - review and editing. SMN: conceptualization; methodology; writing - review and editing; resources; supervision; funding acquisition. FMS: conceptualization; methodology; writing - review and editing; resources; supervision; funding acquisition

## CONFLICT OF INTEREST

The authors declare that they have no conflict of interest.

## DATA AVAILABILITY

Data from this study is available on the Zenodo repository at https://doi.org/10.5281/zenodo.18611037.

## Notes

### Competing Interest Statement

The authors have declared no competing interest.

### Summary of Updates

The text from the introduction and the abstract was slightly modified for grammar and concision matters.

