## supplemental figures for "S6K1 and S6K2 regulate homologous recombination DNA repair through control of BRCA1 protein stability"

### S1. Genotype of S6K1/2 single and double knockouts (SKO and DKO).

**A** SKOs genotype

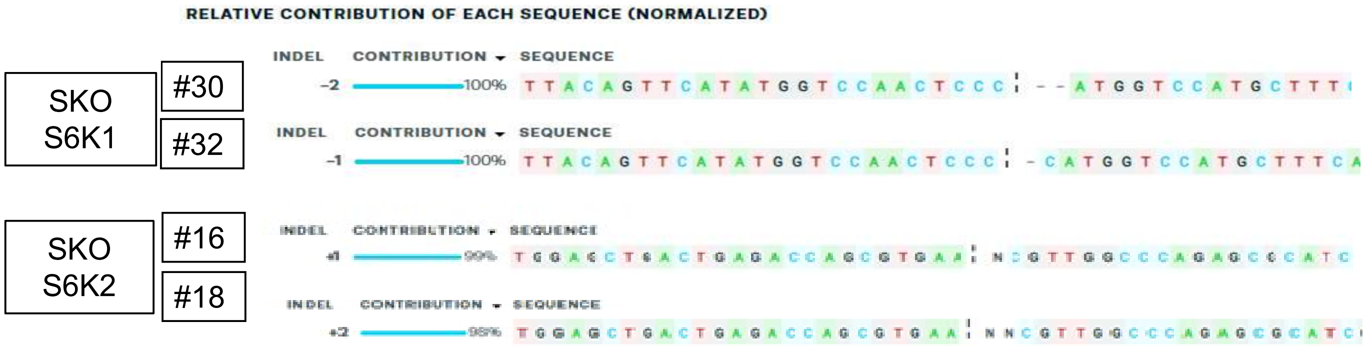


**B** DKOs genotype

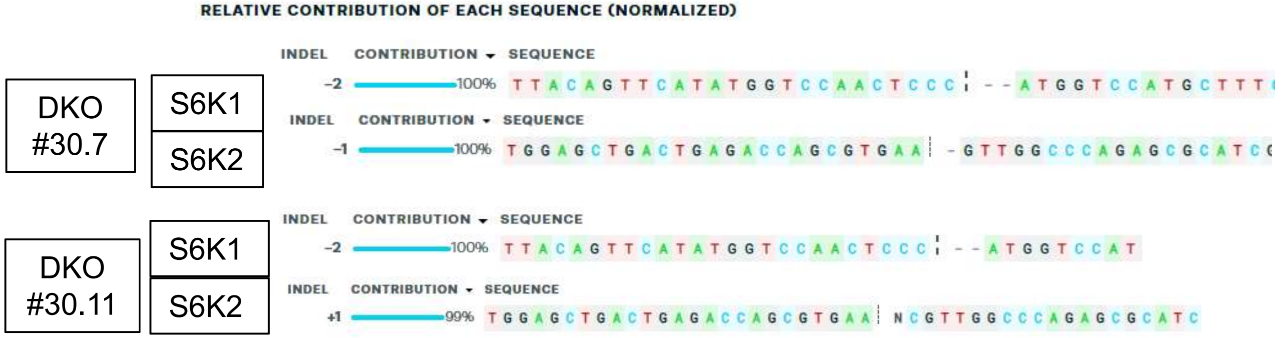

#### S2. Dox-inducible BRCA1 expression vector.

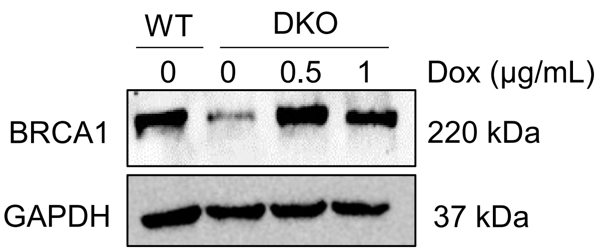


#### S3. Correlation of expression. USP4, mTOR, S6K1 and S6K2 expression.

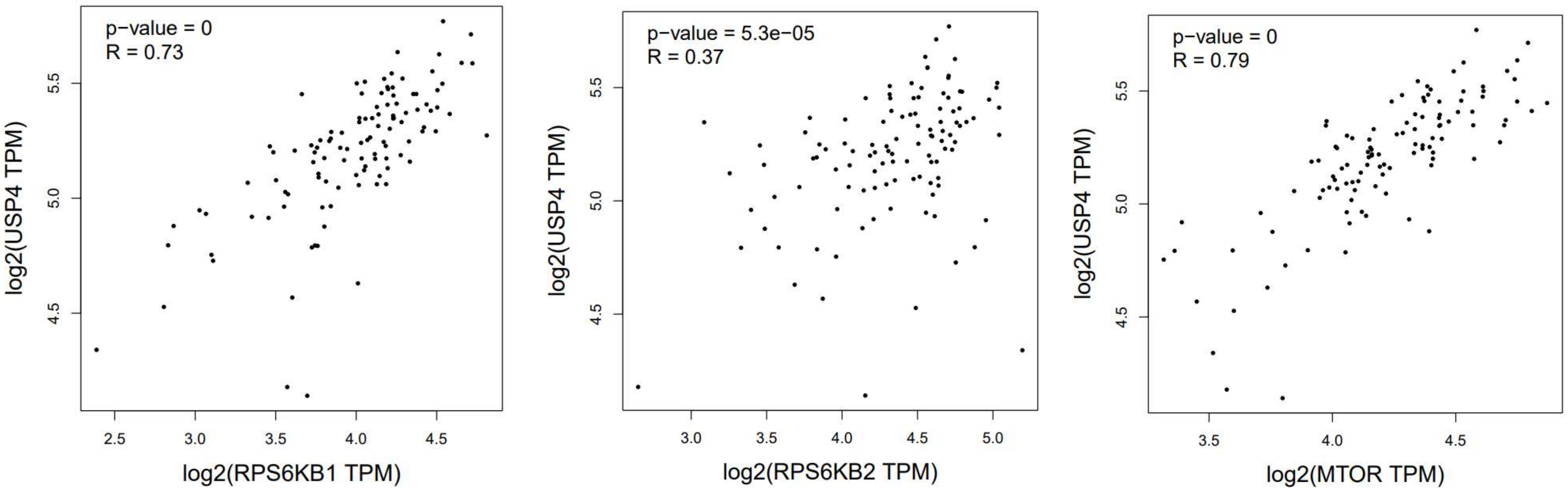
